# Beta bursts question the ruling power for brain-computer interfaces

**DOI:** 10.1101/2023.09.11.557139

**Authors:** Sotirios Papadopoulos, Maciej J Szul, Marco Congedo, James J Bonaiuto, Jérémie Mattout

## Abstract

Current efforts to build reliable brain-computer interfaces (BCI) span multiple axes from hardware, to software, to more sophisticated experimental protocols, and personalized approaches. However, despite these abundant efforts, there is still room for significant improvement. We argue that a rather overlooked direction lies in linking BCI protocols with recent advances in fundamental neuroscience. In light of these advances, and particularly the characterization of the burst-like nature of beta frequency band activity and the diversity of beta bursts, we revisit the role of beta activity in “left vs. right hand” motor imagery tasks. Current decoding approaches for such tasks take advantage of the fact that motor imagery generates time-locked changes in induced power in the sensorimotor cortex, and rely on band-pass filtered power changes or covariance matrices which also describe co-varying power changes in signals recorded from different channels. Although little is known about the dynamics of beta burst activity during motor imagery, we hypothesized that beta bursts should be modulated in a way analogous to their activity during performance of real upper limb movements. We show that classification features based on patterns of beta burst modulations yield decoding results that are equivalent to or better than typically used beta power across multiple open electroencephalography datasets, thus providing insights into the specificity of these bio-markers.

## Introduction

Neural interfaces, and in particular brain-computer interfaces (BCI), have long been conceptualized as effective means of surmounting disabilities for patients suffering from various diseases and traumas, while transhumanist philosophy sees BCI [1] as a way to enhance the capabilities of our bodies and brains. To achieve such goals, a multidisciplinary approach is crucial. Over the past few decades, an increasing number of research groups from diverse fields have been striving towards several objectives, from laying the foundations of BCI [2–6] to improving their reliability [7,8] and applicability under more naturalistic settings [8–10].

Although we are still far from achieving goals like those portrayed in science fiction, a few real-world BCI applications are currently deployed. Most applications revolve around selected groups of patients [12–20], improving their ability to interact with their environment. Such applications usually form part of studies that employ invasive recording techniques in an attempt to acquire high-quality brain signals [21,22]. Invasive techniques provide higher signal-to-noise ratio, spatial specificity and frequency resolution compared to non-invasive techniques, trading off the availability of the subjects, and the necessity of medical interventions. However, the latter attract a significant portion of BCI research due to their safety, the lower equipment cost, and the ability to collect large amount of data from patients and healthy participants. Specifically in the case of electroencephalography (EEG), the added advantage of portability allows for the inclusion of more subjects under more diverse and ecologically valid scenarios, therefore making it currently one of the most attractive platforms.

Non-invasive BCI emerged in the early 90’s [23–25], along with the first spatial filtering algorithms. The Laplacian filter [26,27] allowed for improved signal-to-noise ratio, while the common spatial pattern algorithm (CSP) [28–30] provided a way to weight the contribution of each channel in order to optimize classification. Around the same time, a reliable, reproducible signature of brain activity was demonstrated for the first time, at least on a trial-averaged level. Studies in motor neuroscience involving healthy subjects revealed time-locked changes in induced power within specific frequency bands [31–40]. Brain recordings were shown to exhibit a gradual reduction in signal power, relative to baseline, in the mu (~ 8-12 Hz) and beta (~ 13-30 Hz) frequency bands during an action or during motor imagery (MI): the so-called event-related desynchronization (ERD). This phenomenon is considered to reflect processes related to movement preparation and execution, and is particularly pronounced in the contralateral sensorimotor cortex. Moreover, shortly following the completion of the task, a relative increase in power, the event-related synchronization (ERS), could be observed in the beta band (also referred to as the beta rebound). ERS is thought to reflect the re-establishment of inhibition in the same area.

In the following years, the field witnessed the introduction of more advanced signal processing methods [41], alternative non-invasive recording techniques [42,43] and hybrid BCI paradigms [44–48]. During the past decade, attempts have been made to place more emphasis on the user by studying individual traits that correlate with performance [49], or adapting BCI protocols to the user [50–52] in an effort to better understand and mitigate the problem of BCI illiteracy [8]: the inability of approximately 1/3 of the users to control a motor-imagery based BCI system. Directly linked to this problem, there are significant efforts being made towards creating more informative neurofeedback paradigms by studying the influence of feedback modality [53] and factors not directly linked to the experimental task [54]. This multifaceted endeavor holds the potential of considerably improving existing rehabilitation protocols [55].

Meanwhile, a great body of work has developed an arsenal of advanced pre-processing, feature extraction, and classification algorithms dedicated specifically or adapted to the particular characteristics and limitations of EEG signals [11,56]. As a first step, a standard BCI pipeline includes dimensionality reduction techniques for channel selection and noise removal [57–59]. Subsequently, a common practice for signals recorded during MI or attempted movements is to use a time-frequency (TF) transformation such as the short-time Fourier, Hilbert, or wavelet transform [60–62] and extract the power of the signal in specific time windows and frequency bands of interest. Finally, any of a large range of machine learning algorithms like linear discriminant analysis (LDA) [63–65], support vector machines [66], random forests [67,68] or neural networks [69] can be trained in order to establish a mapping between the features and labels, and assess the performance of the whole pipeline.

This archetypical analysis is, to a significant extent, based on the idea that signal power is the most informative signature of non-invasively recorded neural activity for motor-related tasks. Ever since the characterization of the ERD and ERS phenomena, there has been little to no discussion in the non-invasive BCI field as to whether these features accurately capture the task-related modulations of brain activity. Recent studies in neurophysiology have challenged this view and have demonstrated that the ERD and ERS patterns only emerge as a result of averaging signal power over multiple trials [70,71]. On a single trial level, beta band activity occurs in short, transient events, termed bursts, rather than as sustained oscillations [70–75]. This indicates that the ERD and ERS patterns reflect accumulated, time-varying changes in the burst probability during each trial. Thus, beta bursts may carry more behaviorally relevant information than averaged beta band power. Indeed, studies in humans involving arm movements have established a link between the timing of sensorimotor beta bursts and response times prior to movement, as well as behavioral errors post-movement [71]. Beta burst activity in frontal areas has also been shown to correlate with movement cancellation [73,76,77] and recent studies show that activity at the motor unit level also occurs in a transient manner, which is time-locked to sensorimotor beta bursts [78,79].

Although beta burst rate has been shown to carry significant information, it still comprises a rather simplistic representation of the underlying activity. Every burst can be characterized by a set of TF-based features: the burst peak time and peak frequency, as well as its duration and its span in the frequency axis [80]. In turn, all these descriptors are extracted using a particular time-frequency transformation and constitute simpler representations of the more complex burst waveform that is embedded in the raw signals, and which is characterized by a stereotypical average shape with large variability around it [81]. The waveform features are neglected in standard BCI approaches, because conventional signal processing methods generally presuppose sustained, oscillatory and stationary signals, and are thus inherently unsuitable for analyzing transient activity [82].

In line with the classically described ERD and ERS phenomena, the non-invasive BCI community still heavily relies on signal power as the target feature for classification, although, notably, state of the art Riemannian classifiers [83–85] and some deep learning approaches [86,87] have independently moved on from explicitly using frequency-specific power features. In this article we propose a shift in perspective, by demonstrating how beta band activity during MI tasks is modulated in terms of patterns of distinctly shaped bursts that are better descriptors of transient activity changes.

We have previously argued that analyzing beta burst activity should enable us to gain access to classification features that are at least as sensitive as beta band power [88]. If this hypothesis is valid, then we should be able to test it and verify it using publicly available datasets. Here, we show that this approach allows us to achieve better classification results than those obtained when assessing signal power in binary MI classification tasks, when comparing burst features to signal power from EEG channels C3 and C4. We validate our approach against six open EEG BCI datasets, and provide links between the decoding performance and the modulation of different features considered for classification across datasets and subjects. Although our results obtained by using beta burst features are in most cases inferior to state-of-the-art, namely because our analysis only included two channels and focused solely on the beta frequency band, they are, conversely, superior to those obtained using only beta band power in these channels. This analysis demonstrates the utility of beta burst analysis for BCI and paves the way to improve classification performance in the near future.

## Materials and Methods

### Datasets

We used six open EEG MI datasets: BNCI 214-001 [89], BNCI 2014-004 [90], Cho 2017 [91], MunichMI [92], Weibo 2014 [93] and Zhou 2016 [94], all available through the MOABB project [95]. Briefly, all datasets contain recordings of subjects who were required to perform sustained motor imagery following the appearance of a visual cue on a screen. For our analysis we only considered trials corresponding to the “left hand” or “right hand” classes even if other classes were available in some of the datasets.

### Data pre-processing

For each dataset, recordings were loaded per subject using the MOABB python package (v0.4.6) MotorImagery class, and were filtered with a low pass cutoff of 120 Hz. The low pass cutoff was set to 95 Hz for the Weibo 2014 dataset, because the corresponding sampling frequency of the recordings is 200 Hz. For most of these datasets numerous channels are available, so we defined a subset of channels over the sensorimotor cortex that we deemed relevant for the task and applied pre-processing (Table 1). Then, in this work, we only analyzed data from channels C3 and C4. Each trial was aligned to the cue onset, and the task period was defined as the time between cue onset and the end of the MI task. We used the time window within one second prior to the cue onset as the baseline period (Table 1). In the case of the Cho 2017 and MunichMI datasets we noted the presence of noise at approximately 25 to 30 Hz that interferes with the burst detection step. We therefore included an extra pre-processing step involving a custom implementation of the meegkit python package (v0.1.3, dss_line function) [96] to remove these artifacts. Considering only this subset of sensorimotor channels and all recording periods, we rejected trials using the autoreject python package (0.4.0) [97] (Table 1).

**Table 1.** Attributes of the datasets used in the study.

| Dataset | # Subjects | (# total channels)<br>Channels used for pre-processing | # Total trials<br>(# after trial<br>rejection) | Baseline<br>period (s) | Task<br>period (s) | Post-task<br>period (s) |
| --- | --- | --- | --- | --- | --- | --- |
| BNCI 2014-001 | 9 | (22)<br>"FC3", "FCz", "FC4", "C3", "Cz",<br>"C4", "CP3", "CPz", "CP4" | 288<br>(207 - 287) | -1.0 – 0.0 | 0.0 – 4.0 | 4.0 – 5.5 |
| BNCI 2014-004 | 9 | (3)<br>"C3", "Cz", "C4" | 680 – 760<br>(269 - 621) | -1.0 – 0.0 | 0.0 – 4.5 | 4.5 – 6.5 |
| Cho 2017 | 49 | (64)<br>"FC3", "FCz", "FC4", "C3", "Cz",<br>"C4", "CP3", "CPz", "CP4" | 200 – 240<br>(77 - 240) | -1.0 – 0.0 | 0.0 – 3.0 | 3.0 – 5.0 |
| Munich MI | 10 | (13)<br>"111", "112", "113", "114", "43",<br>"21", "63", "22", "44", "119",<br>"120", "121", "122" | 300<br>(167 - 299) | -1.0 – 0.0 | 0.0 – 7.0 | 7.0 – 9.0 |
| Weibo 2014 | 10 | (60)<br>"FC3", "FCz", "FC4", "C3", "Cz",<br>"C4", "CP3", "CPz", "CP4" | 140 – 160<br>(32 - 160) | -1.0 – 0.0 | 0.0 – 4.0 | 3.0 – 5.0 |
| Zhou 2016 | 4 | (64)<br>"FC3", "FCz", "FC4", "C3", "Cz",<br>"C4", "CP3", "CPz", "CP4" | 290 – 319<br>(167 - 289) | -1.0 – 0.0 | 0.0 – 5.0 | 5.0 – 7.0 |

### Identification of channel-specific beta band and burst detection

Each subject’s data were first transformed in the time-frequency domain from 1 to 43 Hz using the superlets algorithm [98] with a frequency resolution of 0.5 Hz. We selected the superlets algorithm over other more commonly used methods as it allows us to obtain a more optimal tradeoff between temporal and spectral resolution, and because it has been shown to yield better classification results compared to other approaches [99]. Before proceeding with any further analysis we trimmed 200 to 250 ms from the beginning and end of the epoched data in order to exclude any edge effects introduced by the time-frequency transform.

The power spectral density (PSD) of the baseline period was then computed by averaging the resulting TF matrices over the temporal dimension for each trial and channel of a given subject. Based on the distributions of the PSD peaks we attributed the peaks of the power spectra to either the mu (peaks below 15 Hz) or beta (peaks between 15 and 30 Hz) frequency band and proceeded by analyzing activity in the beta band.

Using a previously published iterative, adaptive procedure, we identified bursts within the beta frequency range from the TF matrix, and then extracted their waveforms from the “raw” time series (after low pass filtering as pre-processing) within a fixed time window of 260 ms, centered on the burst peak [100]. Due to inability to parameterize spectra from all datasets we subtracted twice the standard deviation of the TF before fitting each peak as a 2D Gaussian, instead of subtracting the aperiodic activity from the TF matrices [81,101,102], before detecting beta bursts.

### Feature extraction based on patterns of burst rate modulation

Beta burst waveform analysis was performed for each dataset by creating a dictionary of detected bursts across subjects and experimental conditions (“left hand” or “right hand”) (figure 1). This allowed us to create a matrix of burst waveforms by combining all detected bursts per subject, after robust scaling (scikit-learn package [103], v1.0.2). This representation of burst waveforms is suitable for applying a dimensionality reduction technique in order to better understand the variability in the recorded beta burst shapes. For the remaining of the analysis, we only considered channels C3 and C4, or channels 43 and 44 for the MunichMI dataset.

**Figure 1.**
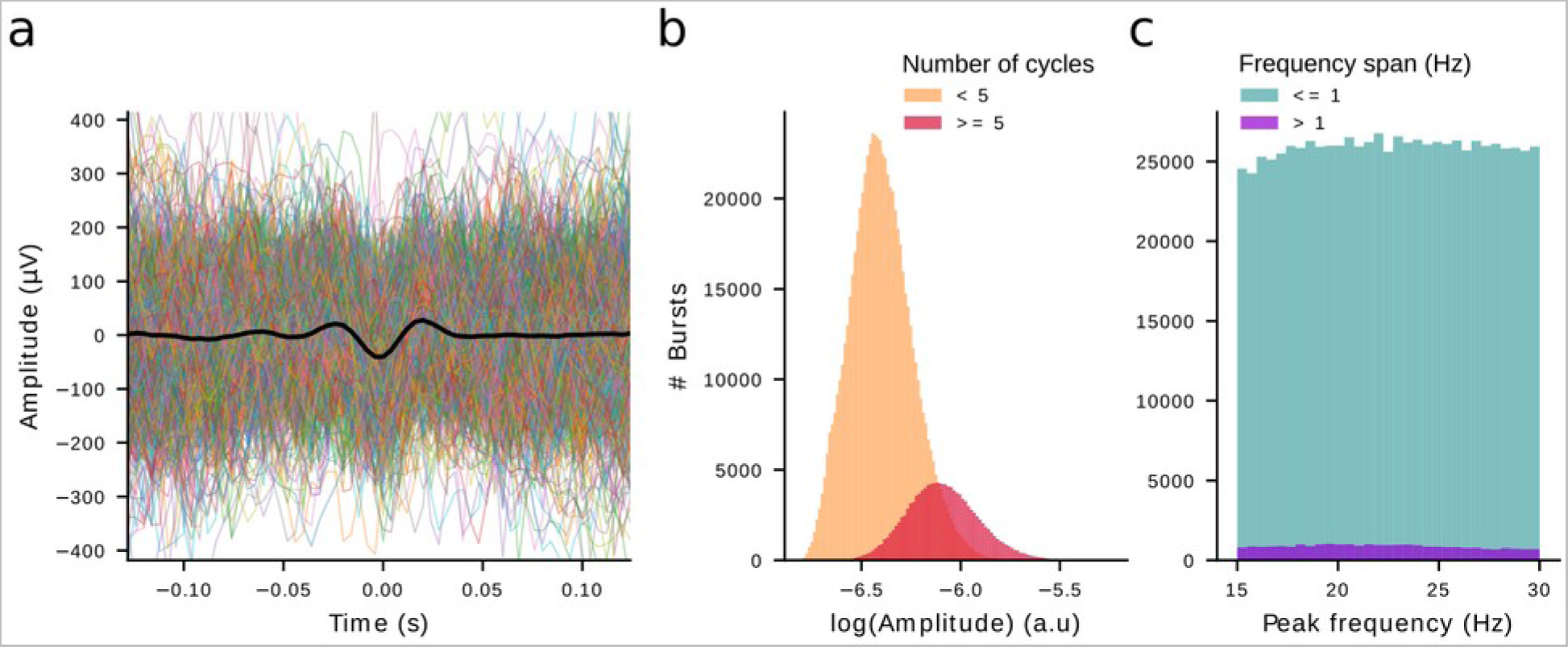
Burst dictionary corresponding to the Zhou 2016 dataset. **(a)** The dictionary contains raw, aligned signal waveforms of 260 ms duration. The black trace represents the average waveform over the whole dictionary. Colored traces correspond to a randomly drawn subset of waveforms (0.2% of all bursts). **(b)** Distribution of the TF amplitude of bursts as computed by the superlets transform, grouped according to burst duration in terms of cycles. The burst detection algorithm identifies a wide range of bursts with amplitudes spanning more than one order of magnitude. The majority of detected beta bursts are low-power, short lasting events. **(c)** Distribution of the peak frequency grouped by the frequency span of each burst. Most of the beta bursts have a narrow frequency span.

Previous work from our group has demonstrated that principal component analysis (PCA) [104] (scikit-learn package, v1.0.2) can be used to understand how the rates of bursts with different waveforms are modulated during reaching movements [100]. In order to construct features suitable for classification, we projected the burst dictionary along each principal component. As such, each burst was associated with a specific score along each dimension of the *C*-dimensional space, representing the distance of the burst’s waveform from the average waveform of all bursts, along this dimension. Because of the scarcity of bursts with extreme scores, we winsorized scores outside of the 2^nd^ and 98^th^ percentile of their distribution. For each component, we then discretized the bursts into groups of bursts within equally spaced score ranges, thus grouping bursts with similar waveforms along that dimension. Since each burst occurs in a specific point in time, following this procedure all bursts were represented in a subspace spanned by the dimensions of scores and time. In other words, for each principal component we generated a representation of burst rate as a function of waveform shape.

### Classification

In order to obtain classification results with our beta burst waveform-based features, we used a stratified, repeated cross-validation approach. For each dataset, we first randomized the trials’ order and stratified the total number of trials of each subject in *M=5* strata. Then, we used half of the trials of one stratum for creating an across-subjects burst dictionary, ran PCA on the resulting waveform matrix and kept track of the rest of the stratum’s trials for cross validating the decoding results. For each subject separately, we then projected the bursts of the remaining four strata (the trials not used during the burst dictionary creation step or for cross validation) along each component and, after averaging the burst rate of each group during the task period, we employed a repeated cross validation with *K*=5 folds. For each fold we repeated this procedure for *100* repetitions by shuffling the order of the features. In order to obtain the results for this analysis, we iterated over a number of possible groups *(from 2 to 9)* and principal components *(from 1 to 8)*. We report the maximum classification score in this hyper-parameter space after cross validating each stratum and averaging across all *M* strata. All steps of the analysis are summarized in a flowchart (figure 2).

**Figure 2.**
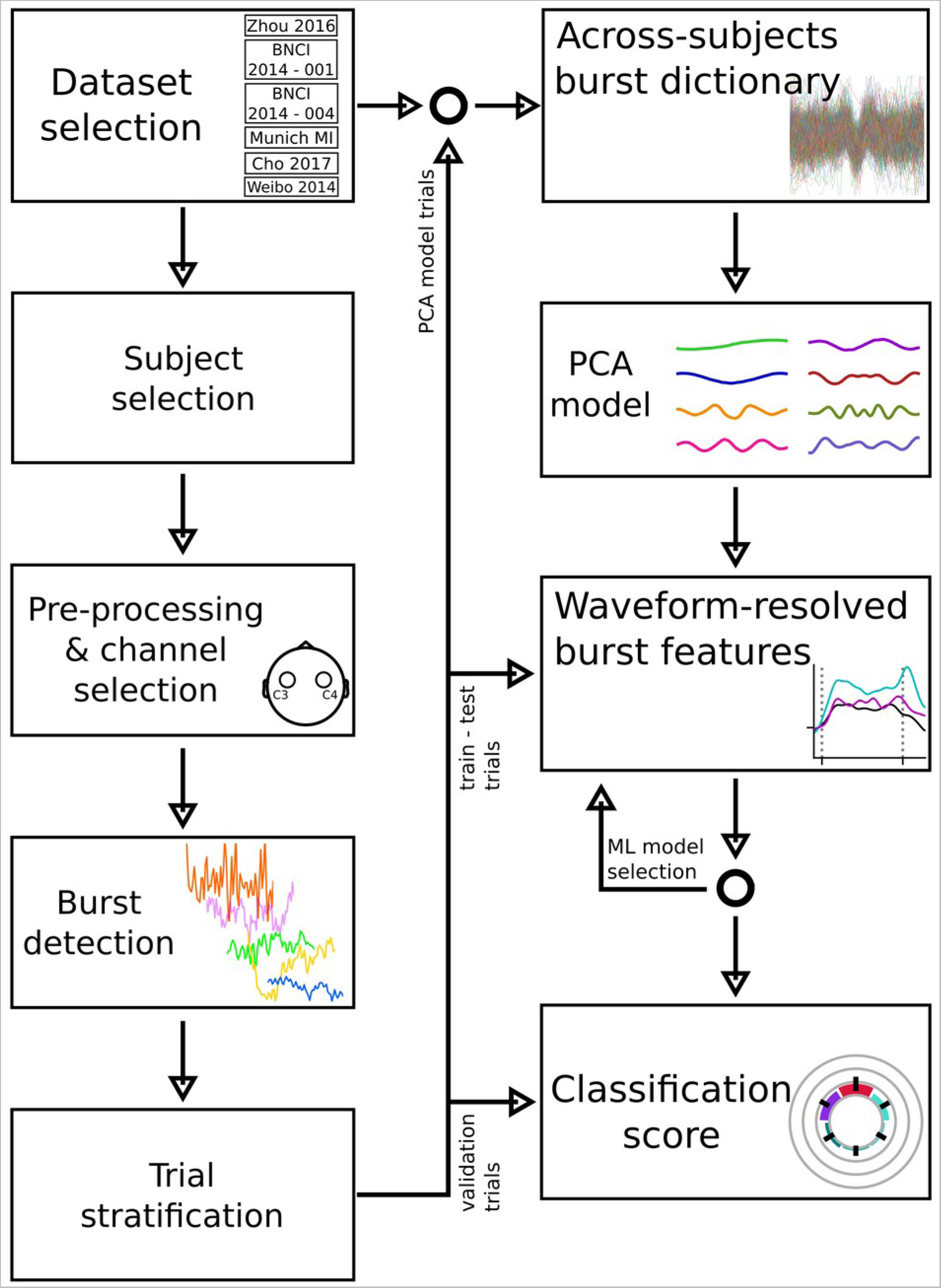
Flowchart illustrating the steps of the proposed analysis. For each dataset, we iteratively pre-processed the data of each subject, rejecting trials and keeping only channels C3 and C4. The burst detection algorithm was run on the raw signals of these two channels. We, then split the remaining trials of each subject in 3 sets. The first set was used only to create the burst dictionary and the corresponding PCA model combining data from all subjects of any given dataset. The second set was used as the training and testing set of trials, in order to select the best model of waveform-resolved features, in terms of decoding score, through a nested, repeated cross validation procedure. Finally, the third set of trials served the role of the validation trials, for the previously selected model.

We compared these results against decoding results obtained by using other related approaches. First, classification results based on beta burst rate were computed for each subject by sampling all detected bursts of channels C3 and C4, and then identifying the rate of bursts within the time course of a trial in non-overlapping time windows of 100 ms. For these results, we only considered bursts with an amplitude equal to or higher than the 75^th^ percentile of the dictionary’s TF amplitude distribution, a threshold commonly used when detecting beta bursts with alternative methods [75,105–108].

We also estimated the decoding accuracy based on TF-based features of the bursts as determined by the burst detection algorithm. We used an approach similar to that described for constructing features and estimating classification results based on burst waveforms. Specifically, for each subject we identified all bursts of channels C3 and C4 and computed the binned burst rate based on the burst volume, burst amplitude, or the combination of TF features, namely burst amplitude, peak frequency, FWHM duration, and FWHM frequency span. We again explored from 2 to 9 possible number of burst groups for each of these features in a repeated, 5-fold cross validation (sup. figure 1).

Band power results for the beta band were based on the power of the Hilbert transform of channels C3 and C4 only. Recordings were first band-pass filtered using the same beta frequency range per channel (15 to 30 Hz). These results are based on a repeated cross-validation approach, and only take into account activity during the task period. The classification features were repeatedly shuffled 100 times, then, for each repetition the trials were split in *K=5* folds.

All classification results were obtained by using LDA as a classifier (scikit-learn, v1.0.2). We estimated the classification score based on the area under the curve (AUC) of the receiver operating characteristic (scikit-learn, v1.0.2). All numeric computations were based on the numpy python package (v1.21.6; [109]), an environment running python (v3.10). We compared trial-level classification results of the waveform-resolved burst features to the beta band power features using a generalized linear mixed model with a binomial distribution and logit link function with correct classification of each trial as the dependent variable, the type of classification feature as a fixed effect, and the subject nested within the dataset as random intercepts. We also compared classification results of the waveform-resolved burst features to the rest of the burst features using a similar model. Statistical analyses were conducted using R (v4.1.2) and lme4 (v1.1-31; [110]). Fixed effects were assessed using type II Wald X^2^ tests using car (v3.1-1; [111]). Pairwise Tukey-corrected follow-up tests were carried out using estimated marginal means from the emmeans package(v.1,8,7 [112]).

## Results

We used six open MI EEG datasets for the purpose of examining the explanatory value of beta burst activity as a feature for BCI classification. For each dataset, we detected beta bursts in a subset of channels over the sensorimotor cortex under two conditions, “left hand” and “right hand” MI. Based on the bursts detected in channels C3 and C4 of each subject, we built dataset-specific burst dictionaries which capture the variability of the burst waveforms (figure 1) (see Materials and Methods).

### Beta bursts with distinct waveforms are characterized by different modulation patterns

We used principal component analysis (PCA) to explain the variability of the burst waveforms within each dictionary (number of components explaining 99% of variance). This method allowed us to reduce the dimensionality of the burst waveform space, with each resulting dimension being a linear combination of the burst waveforms, that emphasizes specific time points that best describe the waveform variability (figure 3 a). Every component defines a motif, along which the waveforms vary. The projection of a burst waveform along each component, associates this waveform with a score, a value that indicates its similarity to the average waveform of bursts within the dictionary along that dimension.

**Figure 3.**
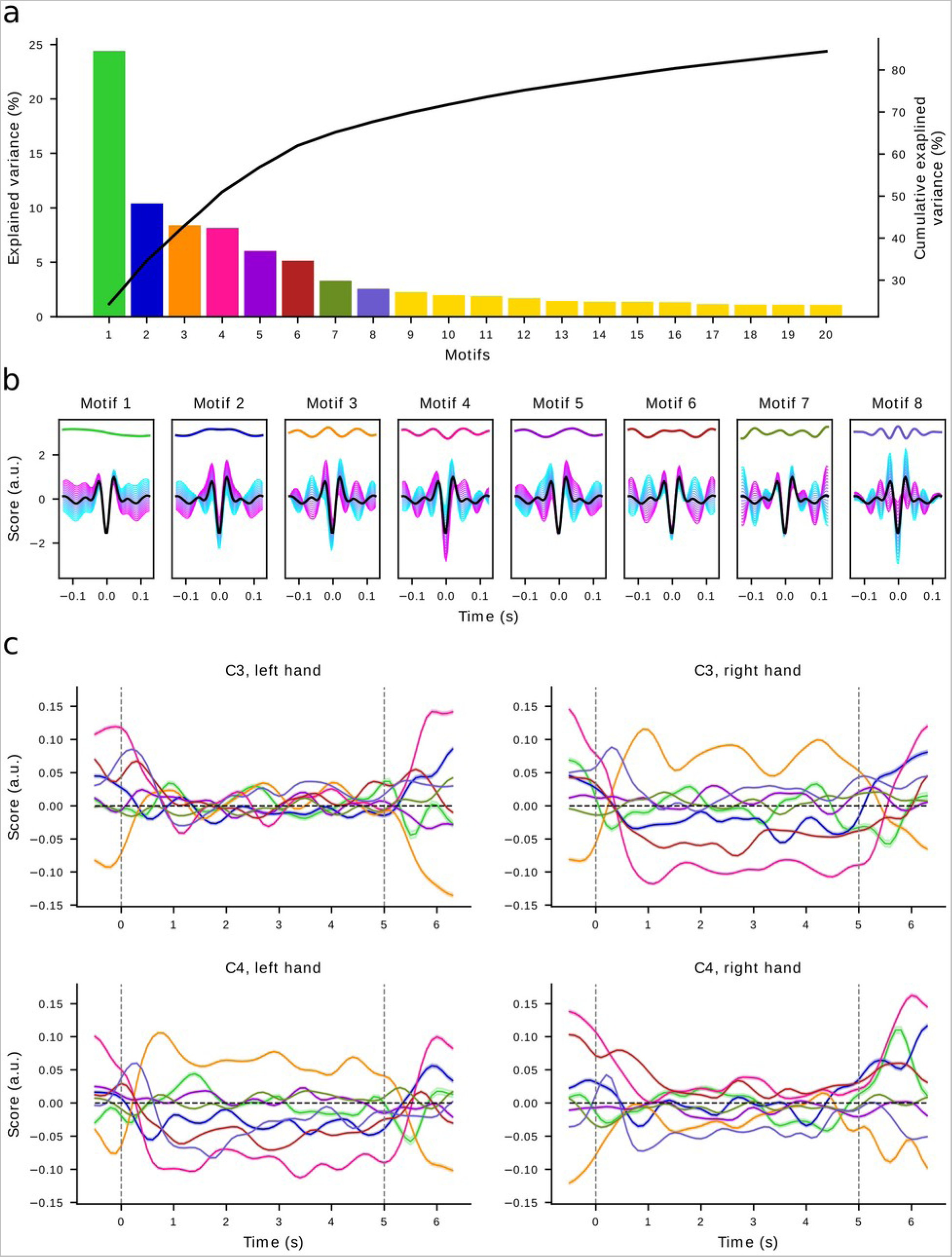
PCA applied on the burst dictionary of the Zhou 2016 dataset. Principal components describe the variability of burst waveforms. **(a)** Ratio of explained variance and cumulative explained variance for the first 20 components. **(b)** The first 8 components define orthogonal axes of waveform shape alteration with respect to the average waveform (black trace). Each subplot depicts one motif (color code as in **a**), the mean waveform (black trace), and simulated waveform alterations along each component, spanning a continuous space from negative (cyan traces) to positive (magenta traces) scores. **(c)** Average score and standard error of all waveforms along each component during the three trial periods for the first 8 components (color code as in **a**) for each condition and channel. During the baseline and post-task periods (signified by the vertical dashed lines), waveforms deviate from the average waveform (score equal to 0) mainly along the third and fourth dimension ipsilaterally, while contralaterally the deviation is more pronounced during the task period.

We simulated how each motif alters the waveform with respect to the average by varying the score along each dimension, adding the weighted eigenvector to the mean waveform (figure 3 b) in order to understand how the burst waveform is modulated by the first 8 motifs. For example, the first motif represents a trend that describes how the waveforms are temporally skewed. Motifs 5, 6 and 7 mainly capture the variability along the flanks of the waveform, whereas motifs 2, 3 and 4 seem to describe changes of the central negative deflection.

For each condition, channel and component we computed the average score of all bursts within the burst dictionary from the baseline to the post-task period, and applied a smoothing kernel of size 2. Burst scores in specific motifs were modulated to different extents within the three trial periods: baseline, task and post-task period (figure 3 c). This means that, on average, bursts with different waveforms occurred more or less frequently within specific trial periods (e.g. motif 4). However, a change in mean waveform shape is ambiguous with respect to the underlying mechanism: e.g. over contralateral motor cortex there was a pronounced decrease in score along component 4 during the task, but this could be due to a reduction in the rate of bursts with high scores, an increase in the rate of bursts with negative scores, or a combination of the two.

To better understand the rate modulation of bursts with distinct waveforms along each component over all experimental periods, we visualized the trial-averaged, baseline-corrected burst rate as a function of time and component score, for the first five components of a representative subject (figure 4; Zhou 2016 dataset, S1). In this particular case there were differences in burst rate modulation between channels C3 and C4, as well as between the two experimental conditions. During the task period there was a decrease in the rate of bursts with large positive or negative scores along component 4 on the contralateral channel for either condition. These patterns correspond to bursts whose waveforms resemble the corresponding magenta and cyan traces. The lateralization of beta burst rate modulation is further exemplified when visualizing the difference between the two channels. The comparison of these differences across the two conditions, reveals that all components and especially components 3, 4 and 5 encode disparities between the “left hand” and “right hand” conditions, and could therefore constitute informative features for a classifier. Interestingly, some components seem to describe a modulation of waveforms during the post-task period, which is particularly evident for either condition in components 1 and 2.

**Figure 4.**
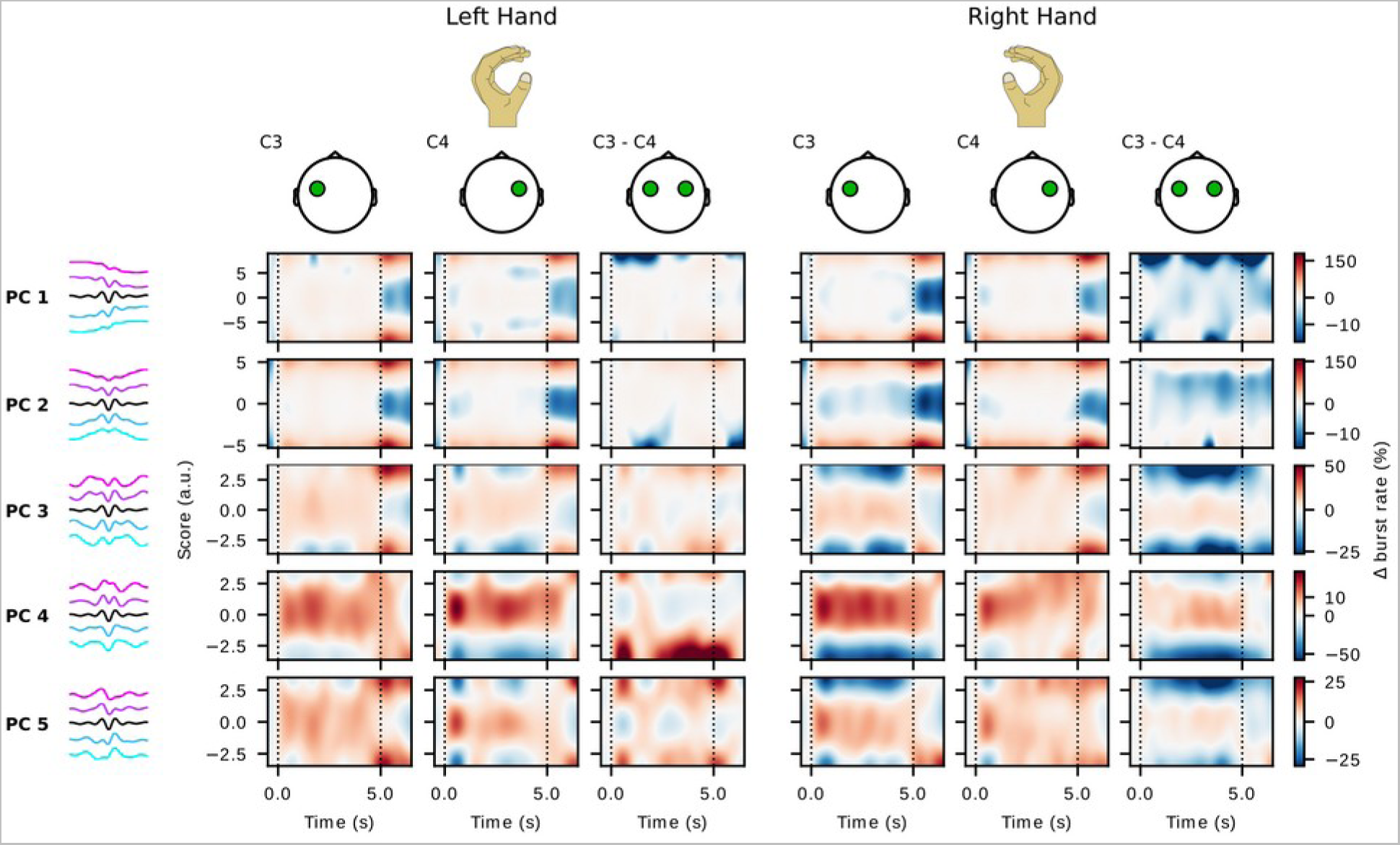
Trial-averaged, baseline-corrected burst rate along different components for a representative subject (Zhou 2016, S1). The first column depicts how burst waveforms vary independently along each component (components as depicted in figure 3). Negative scores correspond to the cyan traces, and positive to the magenta traces. The average waveform is represented by the black trace. During “left hand” trials, burst rate varies per component for channels C3 and C4 and the difference of the two channels. During the task period, both channels exhibit various degrees of burst rate increase for bursts whose waveforms resemble the average along any principal component. Waveforms lying further from the average along component 3 and more prominently 4 are characterized by a reduction of burst rate contralaterally, in channel C4. Similar patterns arise for the “right hand” trials. Component 5 is characterized by an ipsilateral increase and a contralateral decrease of “positive outlier” waveforms. During the post-task period a burst rate increase for specific waveforms is observed, mainly seen along components 1 and 2.

### Beta band burst features outperform beta band power in binary classification tasks

After establishing the lower dimensional space for projecting the burst waveforms, we binned the scores axis into several groups per component (figure 5) using a cross-validation procedure, and analyzed the average burst rate per group (see Materials and Methods). The average burst rate for each group during the task period within each of the two channels was then used as a feature for an LDA classifier, resulting in *G×C×2* features per experimental condition, where G is the number of groups, and C is the number of components, e.g. in the two bottom lines of figure 5 we visualize what would correspond to G=3 and C=2. In order to validate our hypothesis, we compared classification results based on this method against results based on alternative features: the overall beta burst rate for bursts detected in channels C3 and C4 and whose amplitude is greater than a threshold (the 75^th^ percentile of the dictionary’s TF amplitude distribution); time-frequency descriptions of bursts, and band power in the beta frequency (see Materials and Methods).

**Figure 5.**
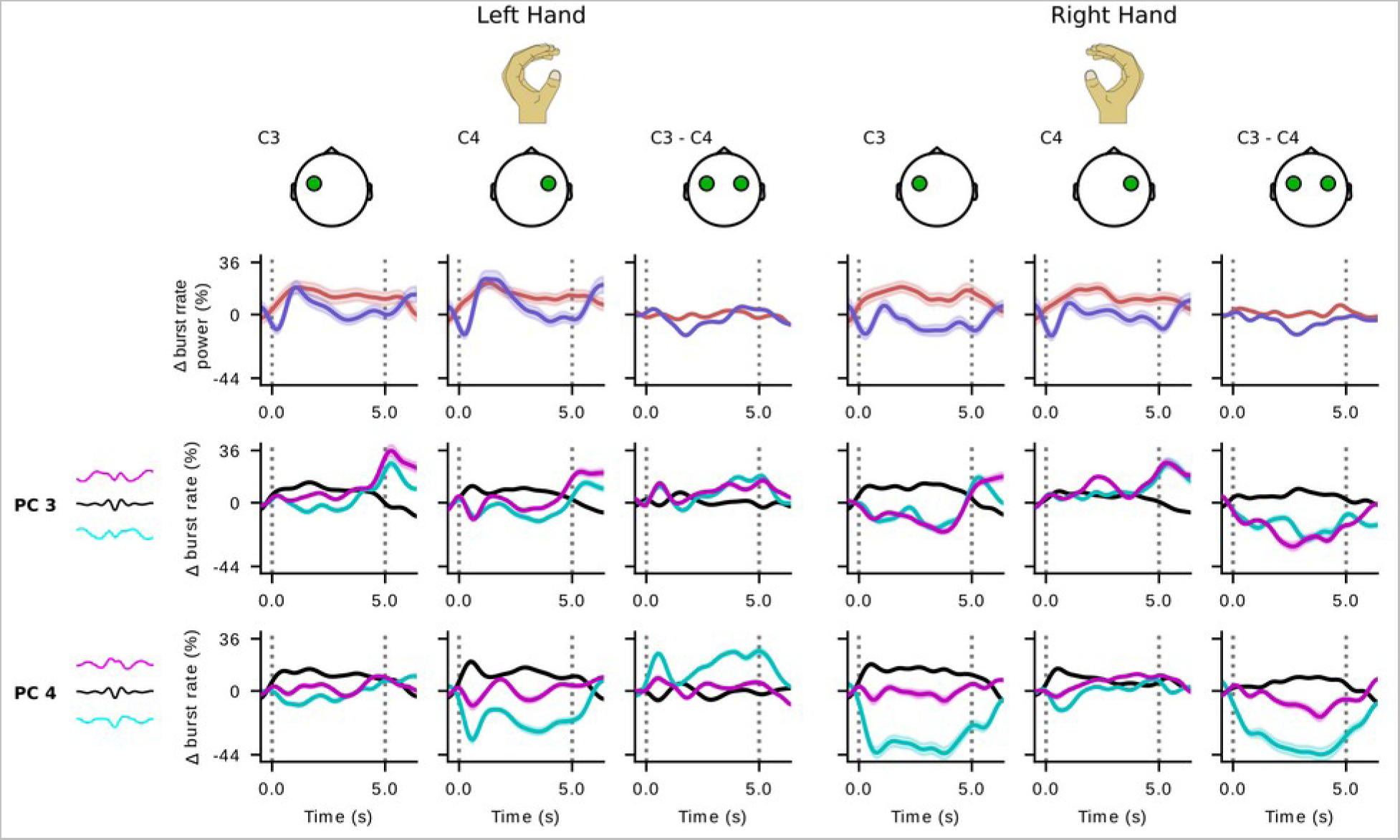
Trial-averaged, baseline-corrected overall burst rate, beta band power and burst rate modulation of three burst groups along components 3 and 4 for a representative subject (Zhou 2016 dataset, S1). For both conditions and channels, beta band power changes (purple trace) roughly track the overall burst rate modulation (red trace). Burst rate modulation for different burst groups varies per condition, channel and component. The differential modulation of burst rate is particularly pronounced contralaterally, in channel C4 during “left hand” trials and channel C3 during “right hand” trials along the fourth component. A clear distinction between conditions is evident when comparing the difference of rate modulation of the two channels for each waveform group.

For each dataset we present the across-subject average results estimated with each method, as well as the results for each participant (figures 6, 7). For the Cho 2017 dataset, which contains a large number of participants, we only show the best ten subjects according to the results based on burst waveform features. The results of all subjects are provided separately (sup. figure 2). At the dataset level, the waveform-resolved burst rate features yield decoding results that are equivalent or better than the results obtained by analyzing beta band power, or alternative beta band representations. These representations appear to bear analogous results in each dataset. We emphasize, though, that the results are highly variable across subjects. For example, for subject S1 of the Zhou 2016 dataset beta power does not hold much explanatory value, unlike beta burst rate, beta burst amplitude or the waveform-resolved burst rate. This is not true for S4 of the BNCI 2014-004 dataset. All representations yield similarly good results, except for the waveform-resolved burst rate that outperforms the rest.

**Figure 6.**
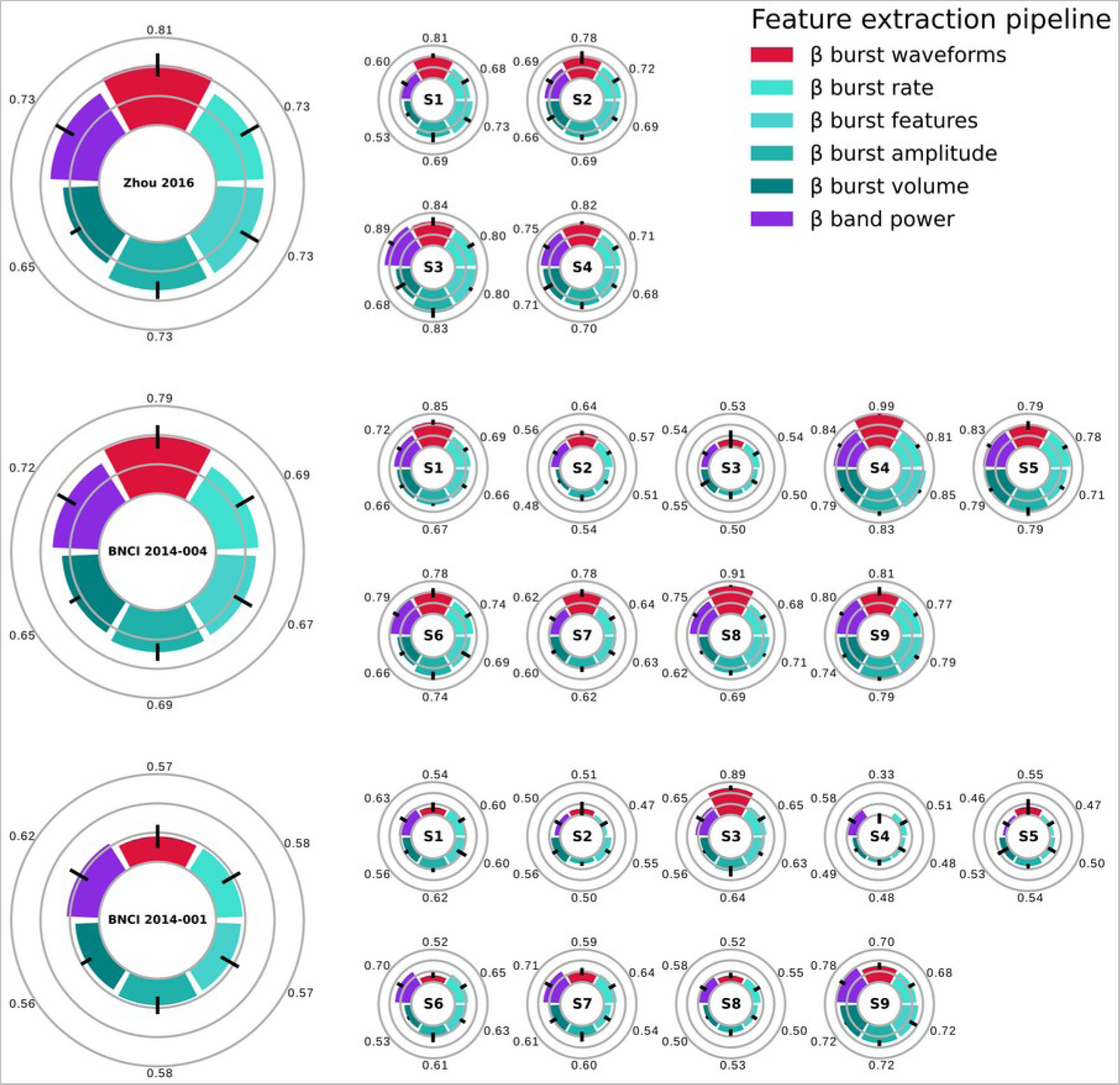
Population average and individual results for binary “left hand” vs “right hand” classification for the BNCI 2014-001, BNCI 2014-004 and Zhou 2016 and datasets. Classification features based on burst waveform-specific rate yield, on average, better results that those obtained using TF-derived burst features, or beta power from channels C3 and C4 across all datasets

**Figure 7.**
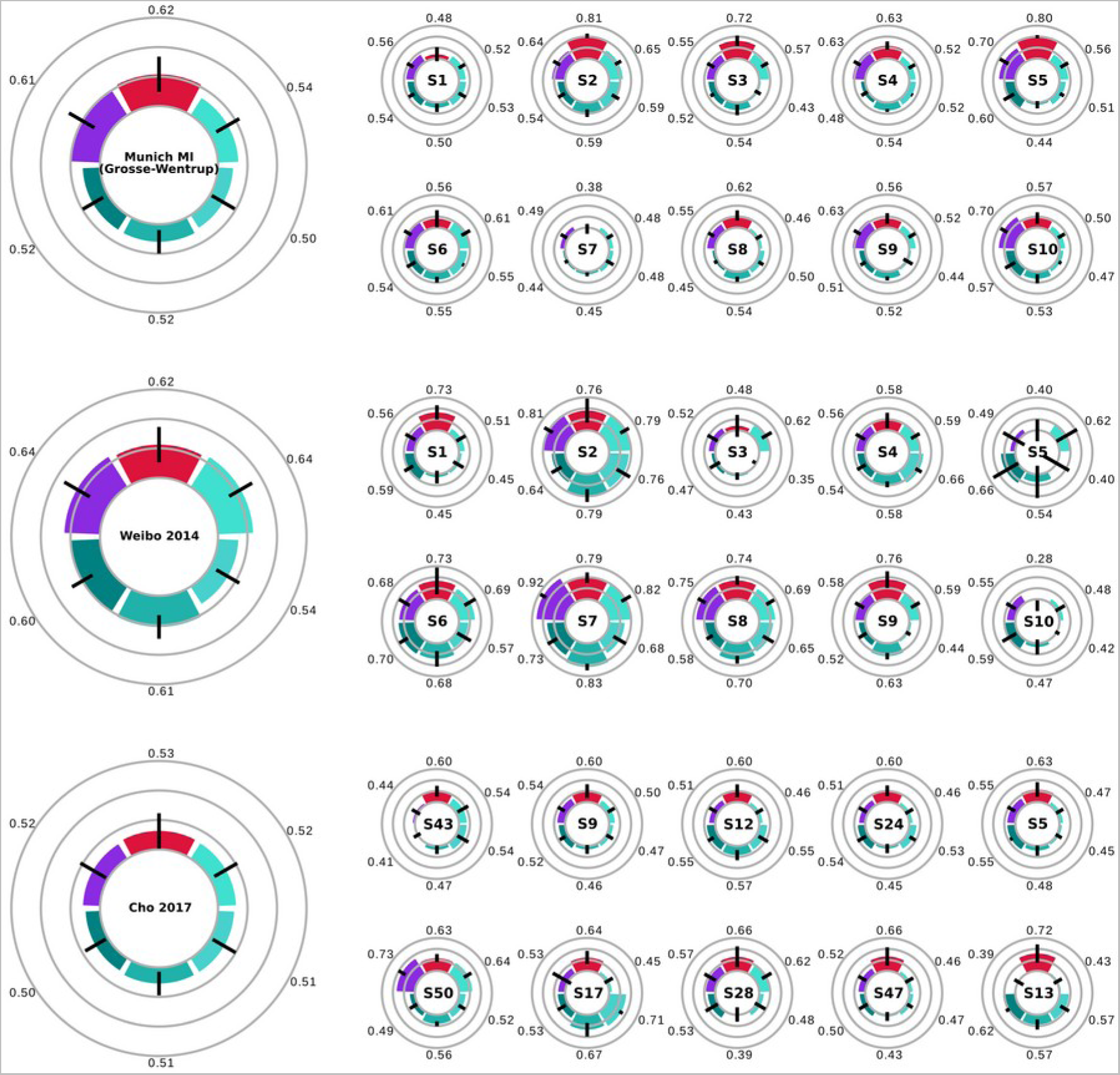
Population average and individual results for binary “left hand” vs “right hand” classification for the Cho 2017, Munich MI (Grosse-Wentrup) and Weibo 2014 datasets. Only the 10 best subjects according to burst waveform features are shown for the Cho 2017 dataset. All features yield equivalent results for the Cho 2017 dataset. Burst waveforms and band power features are equivalent and superior to other beta band activity representations for the Munich Mi dataset. All beta band features except for the combination of multiple features, yield similar results for the Weibo 2014 dataset. Color code as in figure 6.

After obtaining these results we proceeded to quantify the statistical significance of the observed differences for each classification feature set. In order to test the explanatory value of the waveform-resolved burst rate against beta band power we analyzed the decoding results using a generalized linear mixed model (see Materials and Methods). The waveform-resolved burst rate features are significantly better than beta band power features (*X*^2^(1) = 21.384, *p* < 0.001). We also compared the waveform-resolved burst rate against the rest of the examined beta band representations and verified that it yields the highest classification accuracy (*X*^2^(4) = 242.95, all pairwise *p* < 0.001). In conclusion, we confirmed our hypothesis that waveform-resolved beta burst activity holds promise to improve BCI performance, especially if further optimized so that it can be analyzed online and take into account multiple recording channels.

## Discussion

In this study, we showed for the first time that waveform-specific beta burst rate is a representation comparable to beta power within a framework of binary classification MI tasks. In an attempt to understand why, we compared multiple representations of beta activity modulation during the MI task. We showed that bursts of different shapes are selectively modulated following task onset, with distinct waveforms occurring with different probability during different points in time [100] (figures 4 and 5). This modulation can be encoded either by TF-derived features, or alternatively, burst waveforms. All of the TF-derived features were as informative as the overall burst rate when used as classification features, but less reliable than waveform-based features, across all datasets.

The results presented in this article are based on features of beta bursts detected from only two channels, and are therefore not directly comparable to results of previous studies that have implemented standard designs within the BCI literature [95,113] and incorporate all available recording channels, do not perform trial rejection, and utilize spatial filtering. However, waveform-based burst rate features are more informative about imagined movements than beta power in channels C3 and C4. In this regard, our analysis is a first step in the direction of establishing a neurophysiologically informed alternative to currently existing methodologies of feature extraction.

Our results rely on burst dictionaries that combine data from all subjects across a dataset. We have introduced this “transfer learning-like” approach because we have observed that it makes the dimensionality reduction step less susceptible to noise and it results in the same components for all subjects within a dataset, thus rendering the classification features and decoding results easier to interpret. Additionally, it is worth mentioning that due to the enforced orthogonality between the PCA dimensions, the resulting principal components are similar to a Fourier decomposition of the time series, which may be suboptimal by failing to capture components that optimally separate bursts that are differently modulated by the task. Conversely, this property of PCA imposes restrictions on the resulting components that make them similar across datasets (sup. figure 4). This property could be taken advantage of and used in future work for cross-dataset transfer learning.

An important question is whether this procedure would be suitable for online, real-time decoding. The superlets algorithm, and to a lesser extent the burst detection algorithm, are computationally expensive and increasing the number of recording channels, task duration, and frequency resolution would make it difficult to employ this analysis online. However, our results show that beta bursts with particular waveforms are more informative of MI than others. These waveforms could be used as kernels and convolved with online recordings to efficiently detect bursts directly in the time domain. If burst waveforms are maintained across recording sessions, the superlets-based burst analysis could be performed during an offline session and its results used for online burst detection during follow-up, online sessions.

Although we observe distinct patterns of beta burst rate modulations during trials, we do not know how these patterns evolve over sessions and whether or not they are affected by learning. Likewise, how these patterns are influenced by various brain disorders and diseases remains to be studied. There is evidence that beta burst activity is profoundly altered in Parkinson’s disease [75,105,106,114,115], and it could be hypothesized that the alterations in beta band activity following stroke [116–118] may be linked to changes in beta burst waveforms as well. To answer these questions, a longitudinal comparison between a healthy population and clinical patients is needed to establish a link between behavioral or clinical changes and the recorded waveform-specific burst rate patterns or other beta activity representations. Beta burst waveforms could thus serve as an alternative bio-marker for neurofeedback paradigms, and particularly neurorehabilitation protocols.

Tremendous efforts to improve the reliability of non-invasive BCI have been so far unable to provide solutions that would be acceptable for widely-adopted applications. Ever since the characterization of the event-related synchronization and desynchronization phenomena of mu and beta activity, little effort has been put into revisiting the features that are considered to best capture the underlying brain activity in these BCI paradigms. Growing evidence suggests that beta activity modulations are best described in terms of bursts. The analysis presented in this study serves as a proof of concept for the proposed methodology, but there is significant potential for improvement in the burst detection and feature creation procedures. Future directions of interest lie in incorporating more advanced spatial filtering with the burst detection technique, and possibly the use of state-of-the-art Riemannian methods, so that we can leverage the activity of more channels within this framework. Finally, another future direction lies in the incorporation of novel neurophysiological markers for the mu frequency band in our framework. A growing number of studies have shown that the activity in this band can occur as longer-lasting bursts [119], or non-sinusoidal oscillations [120]. We believe that by adapting our approach to the characteristics of this frequency band, or by adopting alternative frameworks such as cycle-by-cycle analysis [121] we can uncover features that will further help us attain the goal of improving BCI robustness. We believe all these goals to be particularly interesting because they hold the promise of further improving current results and rendering them comparable to state-of-the-art approaches.

## Conclusion

Waveform-resolved patterns of burst rate constitute a new way of analyzing beta band activity during motor imagery tasks. The assessment of this method against multiple open EEG datasets shows that this representation is better than conventional power features in terms of classification. This work serves as a first step and opens up numerous directions for further improvements that can potentially ameliorate the reliability of existing, non-invasive brain-computer interface technology.

## Acknowledgments

This work was performed within the framework of the LABEX CORTEX (ANR-11LABX-0042) of Université de Lyon, within the program “Investissements d’Avenir” (decision n° 2019-ANR-LABX-02) operated by the French National Research Agency (ANR). SP, MC, JB, and JM are supported by the French National Research Agency (ANR) project HiFi (2020–2024). MS and JB are supported by the European Research Council (ERC) under the European Union’s Horizon 2020 Research and Innovation Programme (ERC consolidator grant 864550 to JB).

## Data availability Statement

All data are available via the [MOABB project](https://moabb.neurotechx.com/docs/index.html). All scripts necessary for reproducing the results of this article are available at the following public repository: https://gitlab.com/sotpapad/bebopbci.

## Author Contributions

SP, JB and JM conceptualized the manuscript. SP drafted the manuscript and performed the analysis. All authors contributed to manuscript revision, read, and approved the submitted version.

---

**Sup. Figure 1.**
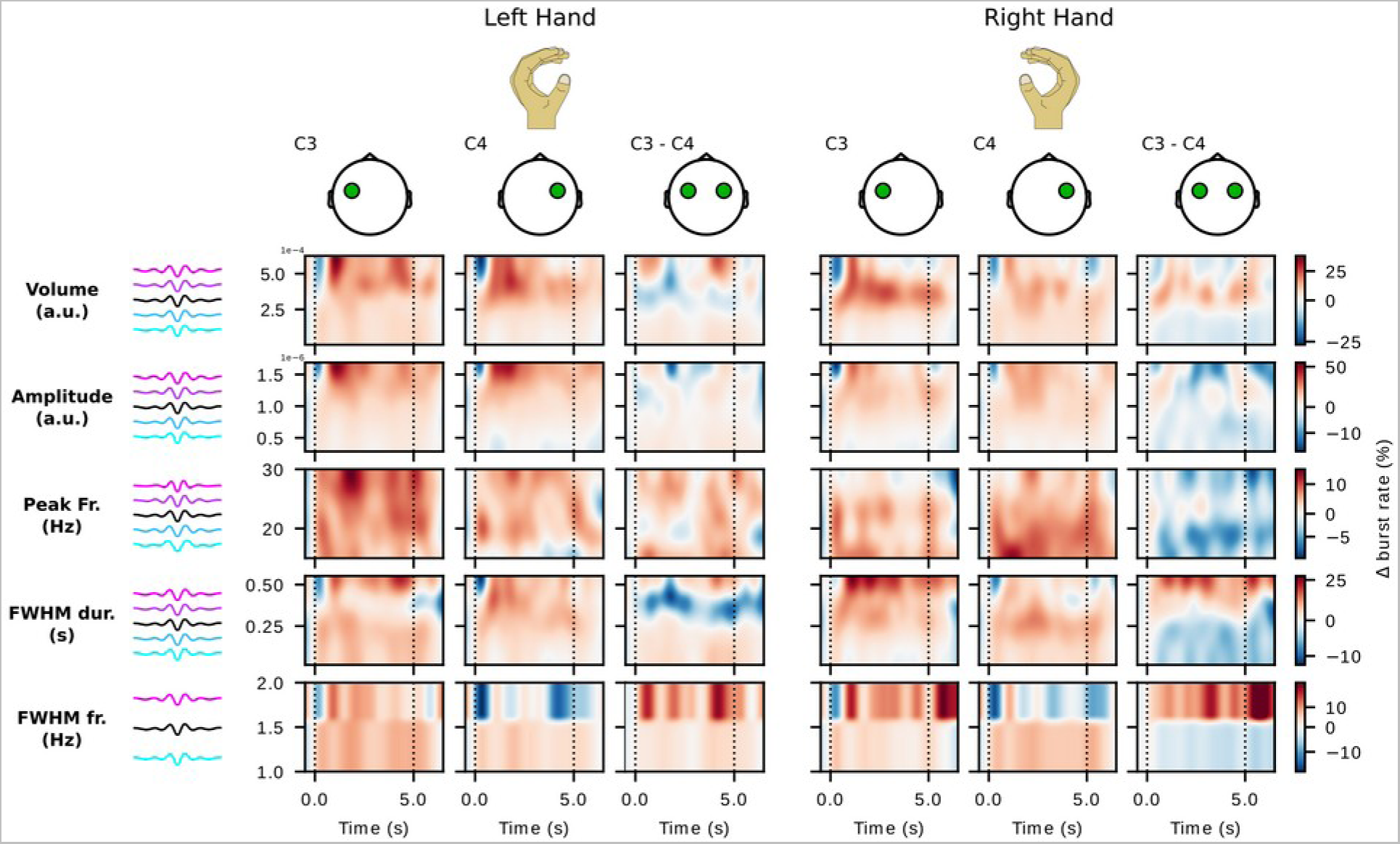
Trial-averaged, baseline-corrected burst rate along different TF-derived features for a representative subject (Zhou 2016 dataset, S1).

**Sup. Figure 2.**
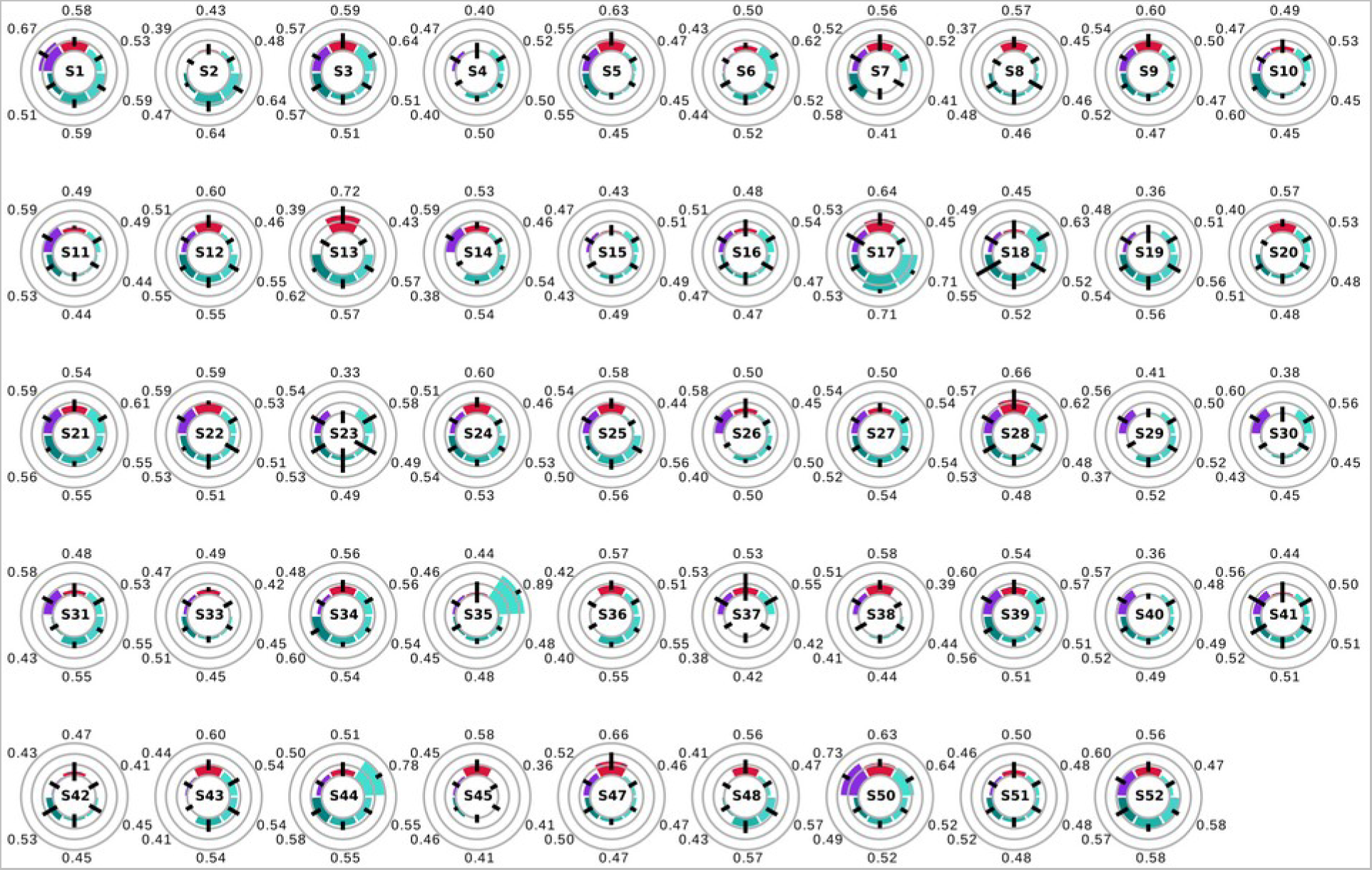
Results for binary “left hand” vs “right hand” classification for all subjects of the Cho 2017 dataset. Color code as in figure 6.

## References

[1] Kurzweil R 2014 The Singularity is Near Ethics and Emerging Technologies ed R L Sandler (London: Palgrave Macmillan UK) pp 393–406

[2] Wolpaw J R 2002 Brain Computer Interfaces for communication and control Front. Neurosci. 4 767–91

[3] Wolpaw J R, Millán J del R and Ramsey N F 2020 Brain-computer interfaces: Definitions and principles Handb. Clin. Neurol. 168 15–23

[4] Ramadan R A and Vasilakos A V. 2017 Brain computer interface: control signals review Neurocomputing 223 26–44

[5] Lotte F, Nam C S and Nijholt A 2018 Introduction : Evolution of Brain-Computer Interfaces Technol. Theor. Adv. Taylor Fr. (CRC Press. 9781498773 1–11

[6] Hatsopoulos N G and Donoghue J P 2009 The Science of Neural Interface Systems Annu. Rev. Neurosci. 32 249–66

[7] Mueller-Putz G, Scherer R, Brunner C, Leeb R and Pfurtscheller G 2008 Better than random: A closer look on BCI results Int. J. Bioelectromagn. 10 52–5

[8] Vidaurre C and Blankertz B 2010 Towards a cure for BCI illiteracy Brain Topogr. 23 194–8

[9] Chavarriaga R, Fried-Oken M, Kleih S, Lotte F and Scherer R 2016 Heading for new shores! Overcoming pitfalls in BCI design Brain-Computer Interfaces 4 60–73

[10] Hughes C, Herrera A, Gaunt R and Collinger J 2020 Bidirectional brain-computer interfaces vol 168 (Elsevier B.V.)

[11] Iturrate I, Chavarriaga R and Millán J del R 2020 General principles of machine learning for brain-computer interfacing Handb. Clin. Neurol. 168 311–28

[12] Blokland Y, Spyrou L, Thijssen D, Eijsvogels T, Colier W, Floor-Westerdijk M, Vlek R, Bruhn J and Farquhar J 2014 Combined EEG-fNIRS decoding of motor attempt and imagery for brain switch control: An offline study in patients with tetraplegia IEEE Trans. Neural Syst. Rehabil. Eng. 22 222–9

[13] Saeedi S, Chavarriaga R and Millan J D R 2017 Long-Term Stable Control of Motor-Imagery BCI by a Locked-In User Through Adaptive Assistance IEEE Trans. Neural Syst. Rehabil. Eng. 25 380–91

[14] Benaroch C, Sadatnejad K, Roc A, Appriou A, Monseigne T, Pramij S, Mladenovic J, Pillette L, Jeunet C and Lotte F 2021 Long-Term BCI Training of a Tetraplegic User: Adaptive Riemannian Classifiers and User Training Front. Hum. Neurosci. 15 1–22

[15] Baniqued P D E, Stanyer E C, Awais M, Alazmani A, Jackson A E, Mon-Williams M A, Mushtaq F and Holt R J 2021 Brain–computer interface robotics for hand rehabilitation after stroke: a systematic review J. Neuroeng. Rehabil. 18 1–25

[16] Luauté J, Morlet D and Mattout J 2015 BCI in patients with disorders of consciousness: Clinical perspectives Ann. Phys. Rehabil. Med. 58 29–34

[17] Mane R, Wu Z and Wang D 2022 Poststroke motor, cognitive and speech rehabilitation with brain-computer interface: A perspective review Stroke Vasc. Neurol. 7 541–9

[18] Chaudhary U, Birbaumer N and Ramos-Murguialday A 2016 Brain–computer interfaces in the completely locked-in state and chronic stroke vol 228 (Elsevier B.V.)

[19] Mcfarland D J 2021 Brain-computer interfaces for amyotrophic lateral sclerosis Dennis Muscle Nerve 61 702–7

[20] Bai Z, Fong K N K, Zhang J J, Chan J and Ting K H 2020 Immediate and long-term effects of BCI-based rehabilitation of the upper extremity after stroke: A systematic review and meta-analysis J. Neuroeng. Rehabil. 17 1–20

[21] Tam W, Wu T, Zhao Q, Keefer E and Yang Z 2019 Human motor decoding from neural signals: a review BMC Biomed. Eng. 1 1–22

[22] Willett F R, Avansino D T, Hochberg L R, Henderson J M and Shenoy K V. 2021 High-performance brain-to-text communication via handwriting Nature 593 249–54

[23] Farwell L A and Donchin E 1988 Talking off the top of your head: toward a mental prosthesis utilizing event-related brain potentials Electroencephalogr. Clin. Neurophysiol. 510–23

[24] Wolpaw J R, McFarland D J, Neat G W and Forneris C A 1991 An EEG-based brain-computer interface for cursor control Electroencephalogr. Clin. Neurophysiol. 78 252–9

[25] Pfurtscheller G, Flotzinger D and Kalcher J 1993 Brain-Computer Interface-a new communication device for handicapped persons J. Microcomput. Appl. 16 293–9

[26] Lu J, McFarland D J and Wolpaw J R 2013 Adaptive laplacian filtering for sensorimotor rhythm-based brain-computer interfaces J. Neural Eng. 10

[27] McFarland D J, McCane L M, David S V. and Wolpaw J R 1997 Spatial filter selection for EEG-based communication Electroencephalogr. Clin. Neurophysiol. 103 386–94

[28] Koles Z J 1991 The quantitative extraction and topographic mapping of the abnormal components in the clinical EEG Electroencephalogr. Clin. Neurophysiol. 79 440–7

[29] Blankertz B, Kawanabe M, Tomioka R, Hohlefeld F U, Nikulin V and Müller K R 2008 Invariant common spatial patterns: Alleviating nonstationarities in Brain-Computer Interfacing Adv. Neural Inf. Process. Syst. 20 - Proc. 2007 Conf. 1–8

[30] Müller-Gerking J, Pfurtscheller G and Flyvbjerg H 1999 Designing optimal spatial filters for single-trial EEG classification in a movement task Clin. Neurophysiol. 110 787–98

[31] Pfurtscheller G and Lopes da Silva F H 1999 Event-relared EEG/MEG synchronization and desynchronization: basic principles Clin. Neurophysiol. 110 1842–57

[32] Pfurtscheller G and Neuper C 1997 Motor imagery activates primary sensorimotor area in humans Neurosci. Lett. 239 65–8

[33] Pfurtscheller G and Berghold A 1989 Patterns of cortical activation during planning of voluntary movement Electroencephalogr. Clin. Neurophysiol. 72 250–8

[34] Pfurtscheller G, Stancák A and Neuper C 1996 Post-movement beta synchronization. A correlate of an idling motor area? Electroencephalogr. Clin. Neurophysiol. 98 281–93

[35] Pfurtscheller G, Brunner C, Schlögl A and Lopes da Silva F H 2006 Mu rhythm (de)synchronization and EEG single-trial classification of different motor imagery tasks Neuroimage 31 153–9

[36] Neuper C, Wörtz M and Pfurtscheller G 2006 Chapter 14 ERD/ERS patterns reflecting sensorimotor activation and deactivation Prog. Brain Res. 159 211–22

[37] Pfurtscheller G, Neuper C, Flotzinger D and Pregenzer M 1997 EEG-based discrimination between imagination of right and left hand movement Electroencephalogr. Clin. Neurophysiol. 103 642–51

[38] Alayrangues J, Torrecillos F, Jahani A and Malfait N 2019 Error-related modulations of the sensorimotor post-movement and foreperiod beta-band activities arise from distinct neural substrates and do not reflect efferent signal processing Neuroimage 184 10–24

[39] Cheyne D and Ferrari P 2013 MEG studies of motor cortex gamma oscillations: Evidence for a gamma “fingerprint” in the brain? Front. Hum. Neurosci. 7 1–7

[40] Kilavik B E, Zaepffel M, Brovelli A, MacKay W A and Riehle A 2013 The ups and downs of beta oscillations in sensorimotor cortex Exp. Neurol. 245 15–26

[41] Makeig S, Enghoff S, Jung T P and Sejnowski T J 2000 A natural basis for efficient brain-actuated control IEEE Trans. Rehabil. Eng. 8 208–11

[42] Waldert S, Preissl H, Demandt E, Braun C, Birbaumer N, Aertsen A and Mehring C 2008 Hand movement direction decoded from MEG and EEG J. Neurosci. 28 1000–8

[43] Naseer N and Hong K S 2015 fNIRS-based brain-computer interfaces: A review Front. Hum. Neurosci. 9 1–15

[44] Allison B Z, Brunner C, Kaiser V, Müller-Putz G R, Neuper C and Pfurtscheller G 2010 Toward a hybrid brain-computer interface based on imagined movement and visual attention J. Neural Eng. 7

[45] Sadeghi S and Maleki A 2018 Recent advances in hybrid brain-computer interface systems: A technological and quantitative review Basic Clin. Neurosci. 9 373–88

[46] Buccino A P, Keles H O and Omurtag A 2016 Hybrid EEG-fNIRS asynchronous brain-computer interface for multiple motor tasks PLoS One 11 1–16

[47] Choi I, Rhiu I, Lee Y, Yun M H and Nam C S 2017 A systematic review of hybrid brain-computer interfaces: Taxonomy and usability perspectives PLoS One 12

[48] Corsi M C, Chavez M, Schwartz D, Hugueville L, Khambhati A N, Bassett D S and De Vico Fallani F 2019 Integrating EEG and MEG Signals to Improve Motor Imagery Classification in Brain-Computer Interface Int. J. Neural Syst. 29 1–12

[49] Lotte F and Rimbert S 2022 How ERD modulations during motor imageries relate to users’ traits and BCI performances 44th International Engineering in Medicine and Biology Conference (Glasgow, United Kingdom)

[50] Lotte F, Jeunet C, Mladenovic J, Kaoua B N and A L P 2018 A BCI challenge for the signal processing community : considering the user in the loop Signal Processing and Machine Learning for Brain-Machine Interfaces pp 1–33

[51] Mladenović J, Frey J, Pramij S, Mattout J and Lotte F 2022 Towards Identifying Optimal Biased Feedback for Various User States and Traits in Motor Imagery BCI IEEE Trans. Biomed. Eng. 69 1101–10

[52] Mladenović J 2021 Standardization of protocol design for user training in EEG-based brain-computer interface J. Neural Eng. 18

[53] Pillette L, N’kaoua B, Sabau R, Glize B and Lotte F 2021 Multi-Session Influence of Two Modalities of Feedback and Their Order of Presentation on MI-BCI User Training Multimodal Technol. Interact. MDPI 5 12

[54] Jeunet C, Lotte F, Batail J M, Philip P and Micoulaud Franchi J A 2018 Using Recent BCI Literature to Deepen our Understanding of Clinical Neurofeedback: A Short Review Neuroscience 378 225–33

[55] Jeunet C, Glize B, McGonigal A, Batail J M and Micoulaud-Franchi J A 2019 Using EEG-based brain computer interface and neurofeedback targeting sensorimotor rhythms to improve motor skills: Theoretical background, applications and prospects Neurophysiol. Clin. 49 125–36

[56] Lotte F, Bougrain L, Cichocki A, Clerc M, Congedo M, Rakotomamonjy A and Yger F 2018 A review of classification algorithms for EEG-based brain-computer interfaces: A 10 year update J. Neural Eng. 15

[57] Zarei R, He J, Siuly S and Zhang Y 2017 A PCA aided cross-covariance scheme for discriminative feature extraction from EEG signals Comput. Methods Programs Biomed. 146 47–57

[58] Kachenoura A, Albera L, Senhadji L and Comon P 2008 ICA: A potential tool for BCI systems IEEE Signal Process. Mag. 25 57–68

[59] Medeiros de Freitas A, Sanchez G, Lecaignard F, Maby E, Barbosa Soares A and Mattout J 2020 EEG artifact correction strategies for online trial-by-trial analysis J. Neural Eng. 17

[60] Bruns A 2004 Fourier-, Hilbert- and wavelet-based signal analysis: Are they really different approaches? J. Neurosci. Methods 137 321–32

[61] Herman P, Prasad G, McGinnity T M and Coyle D 2008 Comparative analysis of spectral approaches to feature extraction for EEG-based motor imagery classification IEEE Trans. Neural Syst. Rehabil. Eng. 16 317–26

[62] Brodu N, Lotte F and Lécuyer A 2011 Comparative study of band-power extraction techniques for Motor Imagery classification IEEE SSCI 2011 - Symp. Ser. Comput. Intell. - CCMB 2011 2011 IEEE Symp. Comput. Intell. Cogn. Algorithms, Mind, Brain 95–100

[63] Pfurtscheller G and Neuper C 2001 Motor imagery direct communication Proc. IEEE 89 1123–34

[64] Vidaurre C, Kawanabe M, Von Bünau P, Blankertz B and Müller K R 2011 Toward unsupervised adaptation of LDA for brain-computer interfaces IEEE Trans. Biomed. Eng. 58 587–97

[65] Llera A, Gomez V and Kappen H J 2014 Adaptive Multiclass Classification for Brain Computer Interfaces Neural Comput. 26 1108–27

[66] Song X, Yoon S C and Perera V 2013 Adaptive Common Spatial Pattern for single-trial EEG classification in multisubject BCI Int. IEEE/EMBS Conf. Neural Eng. NER 19013 411–4

[67] Steyrl D, Scherer R, Oswin F and Gernot R M 2014 Motor Imagery Brain-Computer Interfaces : Random Forests vs Regularized LDA - Non-linear Beats Linear Proc. 6th Int. Brain-Computer Interface Conf. 8–11

[68] Steyrl D, Scherer R, Faller J and Müller-Putz G R 2016 Random forests in non-invasive sensorimotor rhythm brain-computer interfaces: A practical and convenient non-linear classifier Biomed. Tech. 61 77–86

[69] Hazrati M K and Erfanian A 2010 An online EEG-based brain-computer interface for controlling hand grasp using an adaptive probabilistic neural network Med. Eng. Phys. 32 730–9

[70] Jones S R 2016 When brain rhythms aren’t ‘rhythmic’: implication for their mechanisms and meaning Curr. Opin. Neurobiol. 40 72–80

[71] Little S, Bonaiuto J, Barnes G and Bestmann S 2019 Human motor cortical beta bursts relate to movement planning and response errors PLoS Biol. 17 1–30

[72] Lundqvist M, Rose J, Herman P, Brincat S, Buschman T and Miller E 2016 Gamma and beta bursts underlie working memory Neuron 90 152–64

[73] Wessel J R 2020 Β-Bursts Reveal the Trial-To-Trial Dynamics of Movement Initiation and Cancellation J. Neurosci. 40 411–23

[74] Shin H, Law R, Tsutsui S, Moore C I and Jones S R 2017 The rate of transient beta frequency events predicts impaired function across tasks and species Elife

[75] Torrecillos F, Tinkhauser G, Fischer P, Green A L, Aziz T Z, Foltynie T, Limousin P, Zrinzo L, Ashkan K, Brown P and Tan H 2018 Modulation of beta bursts in the subthalamic nucleus predicts motor performance J. Neurosci. 38 8905–17

[76] Hannah R, Muralidharan V, Sundby K K and Aron A R 2020 Temporally-precise disruption of prefrontal cortex informed by the timing of beta bursts impairs human action-stopping Neuroimage 222

[77] Enz N, Ruddy K L, Rueda-Delgado L M and Whelan R 2021 Volume of β-bursts, but not their rate, predicts successful response inhibition J. Neurosci. 41 5069–79

[78] Bräcklein M, Barsakcioglu D Y, Vecchio A Del and Ibáñez J 2022 Reading and Modulating Cortical b Bursts from Motor Unit Spiking Activity 42 3611–21

[79] Echeverria-altuna I, Quinn A J, Woolrich M W, Nobre A C and Ede V 2022 Transient beta activity and cortico-muscular connectivity during sustained motor behaviour Prog. Neurobiol. 102281

[80] Zich C, Quinn A J, Bonaiuto J J, O’Neill G, Mardell L C, Ward N S and Bestmann S 2023 Spatiotemporal organization of human sensorimotor beta burst activity Elife 12:e80160

[81] Szul M J, Papadopoulos S, Alavizadeh S, Daligaut S, Schwartz D, Mattout J and Bonaiuto J J 2023 Diverse beta burst waveform motifs characterize movement-related cortical dynamics Prog. Neurobiol. 165187

[82] Donoghue T, Schaworonkow N and Voytek B 2021 Methodological considerations for studying neural oscillations Eur. J. Neurosci. 1–26

[83] Barachant A, Bonnet S, Congedo M and Jutten C 2012 Multiclass Brain-Computer Interface Classification by Riemannian Geometry IEEE Trans. Biomed. Eng. 59 920–8

[84] Barachant A, Bonnet S, Congedo M and Jutten C 2013 Classification of covariance matrices using a Riemannian-based kernel for BCI applications Neurocomputing 112 172–8

[85] Congedo M, Barachant A and Bhatia R 2017 Riemannian geometry for EEG-based brain-computer interfaces; a primer and a review Brain-Computer Interfaces 4 155–74

[86] Roy Y, Banville H, Albuquerque I, Gramfort A, Falk T H and Faubert J 2019 Deep learning-based electroencephalography analysis: A systematic review J. Neural Eng. 16

[87] Kwon O Y, Lee M H, Guan C and Lee S W 2020 Subject-Independent Brain-Computer Interfaces Based on Deep Convolutional Neural Networks IEEE Trans. Neural Networks Learn. Syst. 31 3839–52

[88] Papadopoulos S, Bonaiuto J and Mattout J 2022 An Impending Paradigm Shift in Motor Imagery Based Brain-Computer Interfaces Front. Neurosci. 15

[89] Tangermann M, Müller K R, Aertsen A, Birbaumer N, Braun C, Brunner C, Leeb R, Mehring C, Miller K J, Müller-Putz G R, Nolte G, Pfurtscheller G, Preissl H, Schalk G, Schlögl A, Vidaurre C, Waldert S and Blankertz B 2012 Review of the BCI competition IV Front. Neurosci. 6 1–31

[90] Leeb R, Lee F, Keinrath C, Scherer R, Bischof H and Pfurtscheller G 2007 Brain-computer communication: motivation, aim, and impact of exploring a virtual apartment. IEEE Trans. neural Syst. Rehabil. Eng. a Publ. IEEE Eng. Med. Biol. Soc. 15 473–82

[91] Cho H, Ahn M, Ahn S, Kwon M and Jun S C 2017 EEG datasets for motor imagery brain-computer interface Gigascience 6 1–8

[92] Grosse-Wentrup M, Liefhold C, Gramann K and Buss M 2009 Beamforming in Noninvasive Brain–Computer Interfaces IEEE Trans. Biomed. Eng. 56 1209–19

[93] Yi W, Qiu S, Wang K, Qi H, Zhang L, Zhou P, He F and Ming D 2014 Evaluation of EEG oscillatory patterns and cognitive process during simple and compound limb motor imagery PLoS One 9 1–19

[94] Zhou B, Wu X, Lv Z, Zhang L and Guo X 2016 A fully automated trial selection method for optimization of motor imagery based Brain-Computer interface PLoS One 11 1–20

[95] Jayaram V and Barachant A 2018 MOABB: Trustworthy algorithm benchmarking for BCIs J. Neural Eng. 15

[96] de Cheveigné A 2020 ZapLine: A simple and effective method to remove power line artifacts Neuroimage 207

[97] Jas M, Engemann D A, Bekhti Y, Raimondo F, Gramfort A, Gramfort A, Automated A and Engemann D A 2017 Autoreject : Automated artifact rejection for MEG and EEG data Neuroimage 159 417–129

[98] Moca V V., Bârzan H, Nagy-Dăbâcan A and Mureșan R C 2021 Time-frequency super-resolution with superlets Nat. Commun. 12 1–18

[99] Bârzan H, Ichim A M, Moca V V and Mureşan R C 2022 Time-Frequency Representations of Brain Oscillations: Which One Is Better? Front. Neuroinform. 16 1–14

[100] Szul M J, Papadopoulos S, Alavizadeh S, Daligaut S, Schwartz D, Mattout J and Bonaiuto J J 2022 Diverse beta burst waveform motifs characterize movement-related cortical dynamics bioRxiv

[101] Brady B and Bardouille T 2022 Periodic/Aperiodic parameterization of transient oscillations (PAPTO)–Implications for healthy ageing Neuroimage 251 118974

[102] Rodriguez-Larios J and Haegens S 2023 Genuine beta bursts in human working memory: controlling for the influence of lower-frequency rhythms bioRxiv 2023.05.26.542448

[103] Pedregosa F, Varoquaux G, Gramfort A, Michel V, Thirion B, Grisel O, Blondel M, Prettenhofer P, Weiss R, Bubourg V, Vanderplas J, Passos A, Cournapeau D, Brucher M, Perrot M and Duchesnay E 2011 Scikit-learn: Machine Learning in Python Fabian J. Mach. Learn. Res. 12 2825–30

[104] Shlens J 2014 A Tutorial on Principal Component Analysis arXiv

[105] Tinkhauser G, Pogosyan A, Little S, Beudel M, Herz D M, Tan H and Brown P 2017 The modulatory effect of adaptive deep brain stimulation on beta bursts in Parkinson’s disease Brain 140 1053–67

[106] Tinkhauser G, Pogosyan A, Tan H, Herz D M, Kühn A A and Brown P 2017 Beta burst dynamics in Parkinson’s disease off and on dopaminergic medication Brain 140 2968–81

[107] Khawaldeh S, Tinkhauser G, Shah S A, Peterman K, Debove I, Khoa Nguyen T A, Nowacki A, Lenard Lachenmayer M, Schuepbach M, Pollo C, Krack P, Woolrich M and Brown P 2020 Subthalamic nucleus activity dynamics and limb movement prediction in Parkinson’s disease Brain 143 582–6

[108] Lofredi R, Neumann W J, Bock A, Horn A, Huebl J, Siegert S, Schneider G H, Krauss J K and Kuühn A A 2018 Dopamine-dependent scaling of subthalamic gamma bursts with movement velocity in patients with Parkinson’s disease Elife 7 1–22

[109] Harris C R, Millman K J, van der Walt S J, Gommers R, Virtanen P, Cournapeau D, Wieser E, Taylor J, Berg S, Smith N J, Kern R, Picus M, Hoyer S, van Kerkwijk M H, Brett M, Haldane A, del Río J F, Wiebe M, Peterson P, Gérard-Marchant P, Sheppard K, Reddy T, Weckesser W, Abbasi H, Gohlke C and Oliphant T E 2020 Array programming with NumPy Nature 585 357–62

[110] Bates D, Mächler M, Bolker B M and Walker S C 2015 Fitting linear mixed-effects models using lme4 J. Stat. Softw. 67

[111] Fox J and Weisberg S 2019 An R Companion to Applied Regression (Sage)

[112] Lenth R V 2023 emmeans: Estimated Marginal Means, aka Least-Squares Means

[113] Luiz P, Rodrigues C, Jutten C and Congedo M 2019 Riemannian Procrustes Analysis : Transfer Learning for Brain-Computer Interfaces IEEE Trans. Biomed. Eng. 66 2390–401

[114] Yeh C H, Al-Fatly B, Kühn A A, Meidahl A C, Tinkhauser G, Tan H and Brown P 2020 Waveform changes with the evolution of beta bursts in the human subthalamic nucleus Clin. Neurophysiol. 131 2086–99

[115] Jackson N, Cole S R, Voytek B and Swann N C 2019 Characteristics of waveform shape in Parkinson’s disease detected with scalp electroencephalography eNeuro 6 1–11

[116] Rossiter H E, Boudrias M H and Ward N S 2014 Do movement-related beta oscillations change after stroke? J. Neurophysiol. 112 2053–8

[117] Shiner C T, Tang H, Johnson B W and McNulty P A 2015 Cortical beta oscillations and motor thresholds differ across the spectrum of post-stroke motor impairment, a preliminary MEG and TMS study Brain Res. 1629 26–37

[118] Kulasingham J P, Brodbeck C, Khan S, Marsh E B and Simon J Z 2022 Bilaterally Reduced Rolandic Beta Band Activity in Minor Stroke Patients Front. Neurol. 13 1–10

[119] Vigué-Guix I and Soto-Faraco S 2022 Using occipital ⍺-bursts to modulate behaviour in real-time bioRxiv

[120] Chen Y Y, Lambert K J M, Madan C R and Singhal A 2021 Mu oscillations and motor imagery performance: A reflection of intra-individual success, not inter-individual ability Hum. Mov. Sci. 78 1–12

[121] Cole S and Voytek B 2019 Cycle-by-cycle analysis of neural oscillations J. Neurophysiol. 122 849–61

